# Cell trajectory modeling identifies a primitive trophoblast state defined by BCAM enrichment

**DOI:** 10.1101/2021.03.27.437349

**Authors:** Matthew Shannon, Jennet Baltayeva, Barbara Castellana, Jasmin Wächter, Samantha Yoon, Jenna Treissman, Hoa T. Le, Pascal M. Lavoie, Alexander G. Beristain

**Author notes:** To whom correspondence should be addressed: Alexander G. Beristain, The British Columbia Children’s Hospital Research Institute, The University of British Columbia, Vancouver, British Columbia, Canada. V5Z 4H4.

## Abstract

In early placental development, progenitor cytotrophoblasts (CTBs) differentiate along one of two cellular trajectories: the villous or extravillous pathways. CTBs committed to the villous pathway fuse with neighboring CTBs to form the outer multinucleated syncytiotrophoblast (SCT), while CTBs committed to the extravillous pathway differentiate into invasive extravillous trophoblasts (EVT). Unfortunately, little is known about the processes controlling human CTB progenitor maintenance and differentiation. To address this, we established a single cell RNA sequencing (scRNA-seq) dataset from first trimester placentas to identify cell states important in trophoblast progenitor establishment, renewal, and differentiation. Multiple distinct trophoblast states were identified, representing progenitor CTBs, column CTBs, SCT precursors, and EVT. Lineage trajectory analysis identified a progenitor origin that was reproduced in human trophoblast stem cell organoids. Heightened expression of basal cell adhesion molecule (*BCAM*) defined this primitive state, where BCAM enrichment or gene silencing resulted in enhanced or diminished organoid growth. Together, this work describes at high-resolution trophoblast heterogeneity within the first trimester, resolves gene networks within human CTB progenitors, and identifies BCAM as a primitive progenitor marker and possible regulator.

**Summary Statement:** Lineage trajectory modeling identifies multiple human progenitor trophoblast states and defines trophoblast differentiation kinetics, where BCAM-expressing progenitors demonstrate enhanced regenerative ability.

## INTRODUCTION

The placenta is a blastocyst-derived tissue that shares the genetic identity of the developing fetus. Derived from cells of the trophectoderm and extraembryonic mesoderm, the human placenta matures into a functional organ by 10-12 weeks’ gestation (GA) (Lee *et al*., 2016; Chang, Wakeland and Parast, 2018). Through its direct interaction with maternal uterine epithelial, stromal, and immune cells, the placenta coordinates the establishment of the maternal-fetal-interface and serves as a critical barrier important for nutrient-waste exchange between maternal and fetal circulations. Trophectoderm-derived trophoblasts perform many functions of the placenta. These include, but are not limited to the facilitation of nutrient and oxygen transfer between mother and fetus (Burton, Cindrova-Davies and Turco, 2020), the modulation of the maternal immune response towards the semi-allogeneic fetus and placenta (Tilburgs *et al*., 2015; Turco and Moffett, 2019), and the production of hormones (i.e. chorionic gonadotropin, progesterone, placental lactogen) and other factors required for pregnancy (Fowden *et al*., 2015).

Of all eutherian mammals, the human placenta is the most invasive. Though humans and rodents both share a haemochorial arrangement of trophoblast and uterine tissue, control of uterine surface epithelial and stromal erosion, as well as artery remodeling by trophoblasts during blastocyst implantation and early placental establishment is far more extensive in humans (Turco and Moffett, 2019). Importantly, key regulators of trophoblast lineage specification in rodents (i.e. *Cdx2, Eomes, Esrrb*, and *Sox2*) do not appear to play essential (or identical in the case of *Cdx2*) roles in human trophoblasts (Knöfler *et al*., 2019). These important differences in human trophoblast biology underlie the need to use human placentas and cell systems for generating fundamental knowledge central to human trophoblast and placental development. While recent advances in regenerative trophoblast culture systems have created the necessary tools required for in-depth cellular and molecular understandings central to trophoblast differentiation (Haider *et al*., 2018; Okae *et al*., 2018; Turco *et al*., 2018), to date, few studies have used these platforms to examine at high resolution trophoblast progenitor and/or stem cell dynamics.

In humans, two major trophoblast differentiation pathways – villous and extravillous – give rise to all trophoblasts in the fetal-maternal interface (Fig. 1A). In the villous pathway, progenitor cytotrophoblasts (CTBs) fuse with neighboring CTBs to replenish or generate new syncytiotrophoblast (SCT), a multinucleated outer layer of the placenta that is the site of transport of most substances across the placenta. Cells committed to differentiate along the extravillous pathway are predicted to originate from CTB progenitors proximal to anchoring villi (Knöfler *et al*., 2019), though populations of villous CTB may already be programmed towards extravillous differentiation or may alternatively harbor a level of bi-potency (Chang, Wakeland and Parast, 2018; Lee *et al*., 2018). Nonetheless, progenitors devoted to this pathway give rise to extravillous trophoblasts (EVTs) that anchor the placenta to the uterine wall (i.e., proximal and distal column CTB). EVTs adopt invasive characteristics hallmarked by interstitial and endovascular EVT subsets, and together facilitate uterine tissue and artery remodeling and modulate maternal immune cell activity (Moffett, Chazara and Colucci, 2017; Pollheimer *et al*., 2018; Papuchova *et al*., 2020). Defects in trophoblast differentiation along either pathway associate with impaired placental function that may contribute to, or drive the development of aberrant conditions of pregnancy (Jauniaux, Moffett and Burton, 2020).

**Figure 1:**
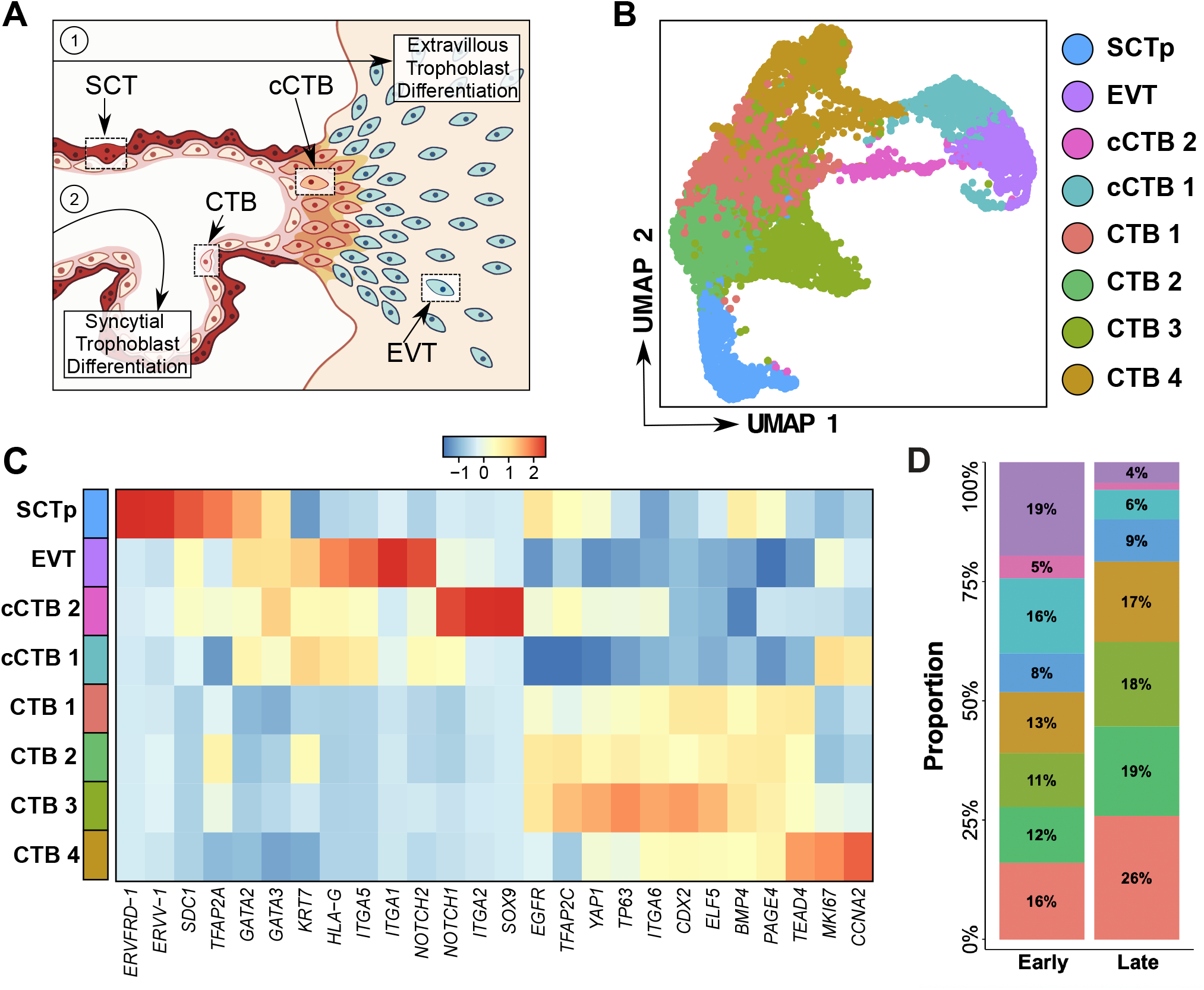
Single cell characterization of first trimester trophoblasts. **(A)** Schematic of a first trimester chorionic villous including anatomical representations of both an anchoring villous and floating villous. Highlighted are trophoblast sub-populations and the villous and extravillous cell differentiation trajectories: cytotrophoblast (CTB), column cytotrophoblast (cCTB), syncytiotrophoblast (SCT), and invasive extravillous trophoblast (EVT). **(B)** Uniform Manifold Approximation and Projection (UMAP) plot of 7798 captured trophoblasts, demonstrating 8 distinct trophoblast clusters: CTB1-4, cCTB1-2, SCTp, and EVT. **(C)** Heatmap showing the average expression of known trophoblast and proliferation gene markers in each trophoblast cluster. Cell cluster names and colours correspond to those in Figure 1C. **(D)** Proportion of early first trimester trophoblasts in each scRNA-seq-defined state in early (6-7 weeks) and late (9-12 weeks) gestational time-points.

In this study, we combined an in-house-generated single cell RNA-sequencing (scRNA-seq) dataset with a publicly available one (Vento-Tormo *et al*., 2018) to define trophoblast heterogeneity and differentiation kinetics in the first trimester of pregnancy. Cell trajectory modeling in chorionic villi and regenerative organoid trophoblasts identified a primitive cell origin. Within this upstream progenitor state, we show elevated expression of the gene basal cell adhesion molecule (*BCAM*), where BCAM enrichment or silencing results in enhanced or impaired trophoblast organoid growth, respectively. Together, this work reaffirms and aligns at high-resolution prior knowledge underlying trophoblast heterogeneity and gene signatures specific to stage of trophoblast development. Further, this work identifies BCAM as a conserved cell-matrix adhesion protein in chorionic villous and trophoblast organoid progenitors important for trophoblast stem cell regeneration and growth.

## RESULTS

### Single cell transcriptomics resolves trophoblast heterogeneity and cell state dynamics

Single-cell transcriptomic analysis enables the identification of novel cell types and states in a systematic and quantitative manner. Here, we applied the 10X Genomics Chromium platform to sequence the human placental transcriptome at single cell resolution (n=7 placentas), and merged this data with single-cell transcriptomic profiles of human first trimester uterine-placental tissues consisting of 4 placentas and 4 decidual specimens (Vento-Tormo *et al*., 2018) (Fig. S1). To examine differences in cellular heterogeneity related to stage of development, we grouped individual samples into early (6-7 weeks’) and late (9-12 weeks’) gestational age (GA) groups. Clinical characteristics and sequencing metrics from each patient and sample are summarized (Table S1). 50,790 cells remained after quality control and data pre-processing, and principal component (PC) analysis validated data integration between the two datasets with separation of cells explained by tissue type, not data source (Fig. S1). Cells clustered into 14 cell types and 31 distinct cell states by transcriptomic similarity, and cell cycle phase proportions per cluster informed the level of proliferation in each cell state (Fig. S1). Decidual cells were identified by vimentin expression and decidual immune cells were specifically identified by *PTPRC* transcripts that encode the leukocyte CD45 antigen. Additional markers were used to identify endothelial, stromal, and fibroblastic cell compartments (Fig. S1).

For trophoblast identification, cells expressing combinations of trophoblast-related transcripts (i.e. *KRT7, EGFR, TEAD4, TP63, TFAP2A, TFAP2C, GATA2, GATA3, HLA-G, ERVFRD-1*), and lacking mesenchymal- and immune-related transcripts (i.e. *VIM, WARS, PTPRC, DCN, CD34, CD14, CD86, CD163, NKG7, KLRD1, HLA-DPA1, HLA-DPB1, HLA-DRA, HLA-DRB1, HLA-DRB5*, and *HLA-DQA1*), were subset for further analysis (Fig. S1). Eight trophoblast clusters from 7,798 cells were resolved (Fig. 1B). Within these 8 clusters, we identified four CTB states (CTB1-4; *TEAD4*^*+*^, *ELF5*^*+*^), a CTB state expressing SCT-associated genes that we termed precursor syncytiotrophoblast (SCTp; *ERVFRD-1*^*+*^, *CGB*^*+*^), two column trophoblast states (cCTB1-2; NOTCH1^+^, ITGA5^+^), and a state specific to EVT (EVT; *HLA-G*^+^, *ITGA1*^+^) (Fig. 1C; Table S2). As expected, CTB1-4 showed varying degrees of enrichment for transcription factors associated with trophoblast progenitor/stemness (i.e., *CDX2, TP63, ELF5, YAP1*), where the CTB3 state showed the greatest level of enrichment of these four genes (Fig. 1C).

Column trophoblasts are precursors to EVT, where cells situated within proximal and distal sites of anchoring columns can be molecularly distinguished by expression of NOTCH and integrin genes (Zhou *et al*., 1997; Haider *et al*., 2016). To this end, cCTB2 expresses high levels of *NOTCH1, SOX9*, and *ITGA2*, the latter of which defines a distinct EVT progenitor population and likely represents a proximal column progenitor (Lee *et al*., 2018) (Fig. 1C). Compared to cCTB2, cells within the cCTB1 state showed higher levels of *ITGA5, NOTCH2*, and *HLA-G*, indicating that this state is developmentally downstream of cCTB2 and cells in this state likely reside within central and distal regions of anchoring columns (Haider *et al*., 2014) (Fig. 1C). Stratifying cells by phases of the cell cycle showed that cCTB1 column cells are predominately in G_2_/M and S phases, while the majority of cCTB2 column cells are in G_1_ (Fig. S1). Terminally differentiated EVT and SCTp states are largely G_1_ cells, as are progenitor CTB1-3 states (Fig S1). This was in contrast to CTB4, where the vast majority of cells are undergoing DNA synthesis and mitosis (Fig. S1), and express hallmark genes of proliferation (i.e. *MKI67, CCNA2*), and show enrichment for the transcription factor *TEAD4* (Fig. 1C). Importantly, these distinct trophoblast states largely align with recent single cell transcriptomic characterizations of cells from first trimester pregnancies (Liu *et al*., 2018; Suryawanshi *et al*., 2018; Vento-Tormo *et al*., 2018).

Having defined eight trophoblast types/states within first trimester placental tissue, we next examined cell state stability across the first trimester. Relative frequencies of CTB1-4 states increase, whereas proportions of cCTB and EVT states decrease from early (6-7 weeks’) to late (9-12 weeks’) gestation (Fig. 1D). Notably, an almost 5-fold reduction in the proportion of EVT (19% to 4%) is observed (Fig. 1D), suggesting that villous development and branching contribute to the increase in CTB frequency. Unexpectedly, and unlike CTB1-4, frequencies in SCTp remained relatively constant (8-9%) (Fig. 1D), though, as with all trophoblast proportions, accuracy is highly dependent on minimizing biases in single cell dissociation and cell capture pipelines prior to barcoding and sequencing. Together, these findings define unique states of trophoblasts in the first trimester placenta and highlight how proportions of these states change between the early and late first trimester, a developmental window during which the human placenta becomes fully functional.

### Single cell trajectory modeling identifies a CTB origin

RNA velocity, the time derivative of the gene expression state, can be used to predict the future transcriptional state of a cell. RNA velocity analysis in combination with single-cell trajectory analyses provides us with insights into the transcriptional dynamics of developing cells and enables high-resolution lineage pathway ordering. By applying scVelo to our dataset to refine progenitor and cell end-point relationships, we identified an origin that overlapped with CTB2; this predicted origin was also proximal to the CTB3 state, indicating that these two states contain a proportionally high content of primitive trophoblasts (Fig. 2A). Two paths were identified extending from this origin towards the EVT state and one path developing from the origin towards the SCTp state. As well, we found an additional differentiation trajectory, extending from CTB2 into CTB3, representing possible CTB progenitor reserves or progenitor states in transition (Fig. 2A). When shown in a 3D perspective, a linkage between CTB3 and CTB4 becomes evident that is not obvious in 2D clustering, indicating that the CTB2 state can transition to the proliferative CTB4 state via multiple cell pathways (Fig. S2). Monocle3 pseudotime ordering was also applied to increase confidence in trophoblast hierarchical ordering. Similar to RNA velocity, CTB2 was identified as a state aligning to the origin (though CTB1 also showed partial overlap); increased complexity is observed as CTBs progress along one of two trajectories that lead to either EVT or SCT end-point states (Fig. S3). Specifically, Monocle3 ordering revealed a looped trajectory within CTB2 that exits towards CTB3, SCTp, and EVT states (Fig. S3).

**Figure 2:**
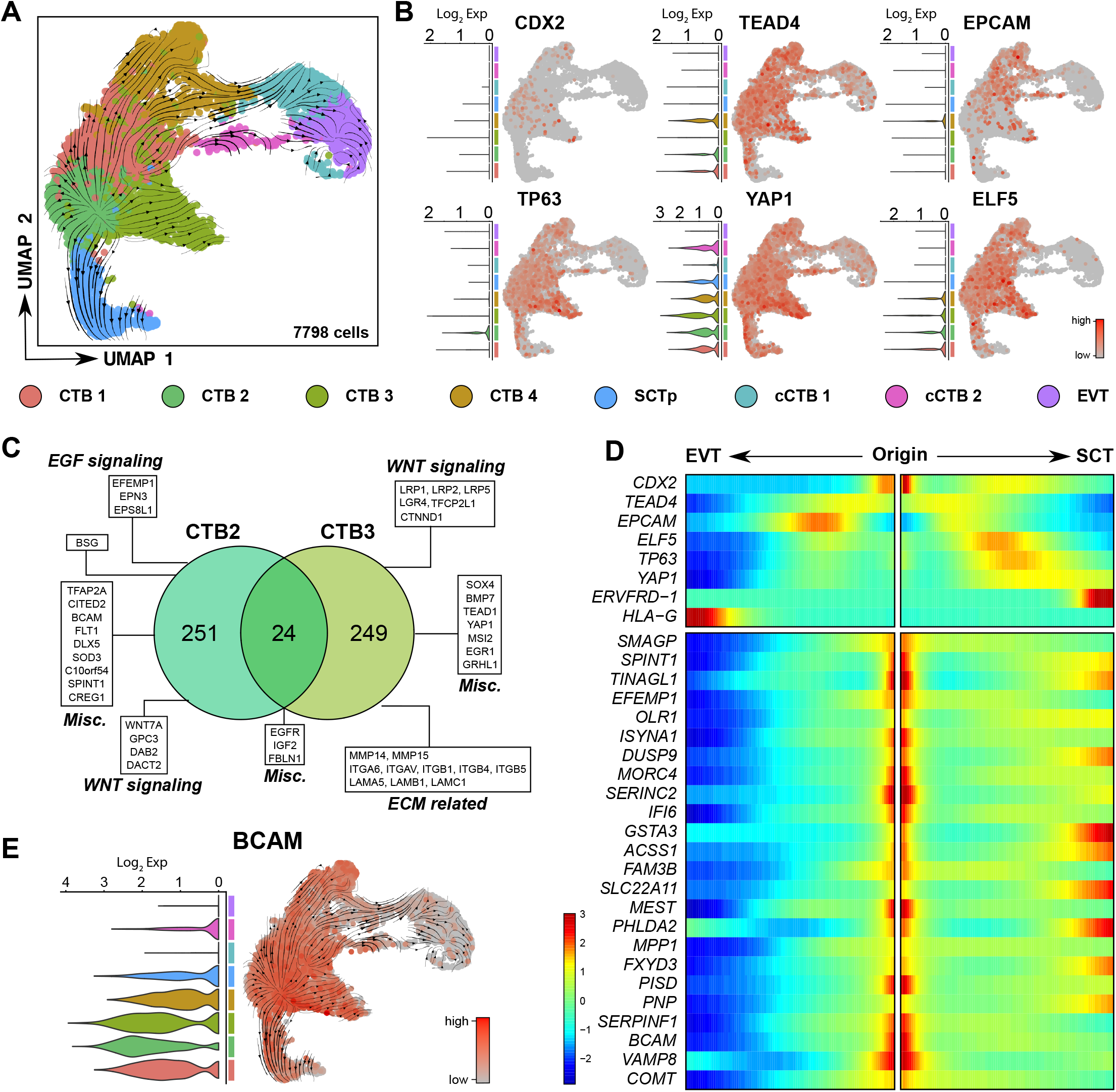
scVelo-informed cell differentiation identifies a primitive CTB origin. **(A)** UMAP plot overlain with projected RNA velocity (scVelo) vector field, indicating trophoblast directionality (represented by the velocity arrows) away from a putative origin towards endpoint SCTp and EVT states. Cell cluster names and colors correspond to those in Figure 1C. **(B)** UMAP plots showing gene expression (red) of progenitor- and stem cell-associated genes in all scRNA-seq-informed trophoblast states. **(C)** Venn diagram comparing elevated genes within CTB states 2 and 3. Highlighted are genes related to WNT and EGFR signaling pathways, genes involved in extracellular matrix (ECM) interactions, and miscellaneous (Misc) genes of interest. **(D)** A heatmap was constructed using inferred pseudotime and the top 23 up-regulated genes in CTB2. Included within the heatmap are known genes that align with the CTB (CDX2, TEAD4, EPCAM, TP63, YAP1, ELF5), the SCT (ERVFRD-1) and EVT (HLA-G) states. **(E)** scVelo-overlaid UMAP and violin plots showing *BCAM* levels in all first trimester trophoblast states.

We next compared these refined trajectories with expression patterns of known CTB progenitor/stem cell genes *CDX2, TEAD4, YAP1, TP63, ELF5*, and *EPCAM*. Predominant expression of *ELF5, TEAD4*, and *YAP1* was observed in all CTB states, while expression of the CTB stem cell transcription factor *CDX2* showed predominant levels in cells in CTB1 and CTB3, and to a lesser extent CTB2 and CTB4 (Fig. 2B). Average *TP63* expression was highest in CTB3, though a greater proportion of cells within CTB2 expressed *TP63* compared to other CTB states, and *EPCAM* showed greatest enrichment within the CTB1 and CTB4 (Fig. 2B). Because the scVelo-predicted origin aligned closely to CTB2 and CTB3, we examined genes both shared and enriched in each of these states (Fig. 2C). Twenty-four genes were shown to overlap between CTB2 and CTB3, including *EGFR, IGF2*, and *FBLN1* (Fig. 2C; Table S3). 249 genes showed enrichment in CTB3, and of these many associate with trophoblast stem cell and regenerative properties (i.e., *YAP1, TP63*), Wnt signaling (*LRP5*), and cell-matrix interactions (*MMP14, MMP15, ITGA6, LAMA5, LAMB1*) (Fig. 2C). The CTB2 state showed enrichment for 251 genes, including the Wnt ligand, *WNT7A*, the laminin receptor basal cell adhesion molecule (*BCAM*), and the transcriptional modulators *TFAP2A* and *CITED2* (Fig. 2C). Expression levels of the top twenty-five CTB2-enriched genes (FDR ranked) plotted by cell state over a simple branched pseudotime [as performed previously using Monocle 2 in (Vento-Tormo *et al*., 2018; Treissman *et al*., 2020)] showed consistent enrichment of all genes to the CTB origin (Fig. 2D). Notably, some genes (e.g., *BCAM, IFI6*) show a consistent down-regulation along the SCT and EVT trajectories, while others (e.g., *DUSP9, GSTA3, VAMP8*) show transient decreases followed by increases in gene levels approaching SCT or EVT trajectory endpoints (Fig. 2D). scVelo-alignment of *BCAM*, a gene encoding a cell-ECM laminin receptor and one of the most enriched genes within the CTB2 state (Table S2), shows that *BCAM* levels are highest within the predicted CTB origin, and gradually decrease as CTBs differentiate into SCTp or cCTB/EVT (Fig. 2E). Taken together, we have identified a primitive CTB state within the first trimester placenta that shows molecular features consistent with that of putative CTB stem cells. We also identify multiple genes aligning with this origin, many of which have not being described in trophoblast progenitor biology, including the gene *BCAM*.

### Trophoblast stem cells recapitulate scRNA-seq-informed CTB heterogeneity and hierarchy

Recently described regenerative human trophoblast models provide an improved method for studying trophoblast differentiation *in vitro* (Okae *et al*., 2018). Three-dimensional organoid cultures established from human trophoblast stem cell lines (hTSCs) also reproduce molecular features underlying CTB maintenance and differentiation (Saha *et al*., 2020). To examine the reproducibility of trophoblast-intrinsic differentiation programs defined by scRNA-seq analysis of chorionic villous tissues, we subjected three hTSC lines to scRNA-seq analysis prior to and following EVT differentiation. hTSC organoids (CT27, CT29, and CT30 CTB lines) were cultured for 7 days in regenerative Wnt^+^ (containing the Wnt potentiators R-spondin and CHIR99021) and EVT-differentiating Wnt^-^ (lacking R-spondin and CHIR99021) media, respectively, prior to single cell workflow. To confirm the intactness of our system, a separate cohort of hTSC organoids were cultured over 21 days in these media and differentiation was assessed by light microscopy, flow cytometry, and qPCR analysis. hTSC-derived organoids cultured in regenerative media established large spherical colonies characterized in part by elevated expression of the CTB progenitor genes *TEAD4* and *ITGA6* (Fig. 3A, 3B). Moreover, expression of surface MHC-I (HLA-A, -B, -C) by flow cytometry did not detect these MHC-I ligands in undifferentiated cultures (Fig. 3C). Comparatively, hTSC organoids cultured in EVT differentiating Wnt^-^ media showed characteristic cellular extensions from the organoid body (Fig. 3A), as well as a marked decrease in levels of *TEAD4* mRNA (Fig. 3B). As expected, both immature (*ITGA5, NOTCH1*) and mature (*NOTCH2, HLA-G*) EVT marker genes, as well as HLA-C and HLA-G surface expression increased along the EVT differentiation timeline (Fig. 3B, 3C). Syncytial formation occurs in both regenerative Wnt^+^ and EVT Wnt^-^ conditions, and this was reflected at the gene level were *CGB* and *CSH1* levels markedly increased over 21 days (Fig 3C). Together, these data indicate that the 3D organoid model to study CTB regeneration and differentiation is intact.

**Figure 3:**
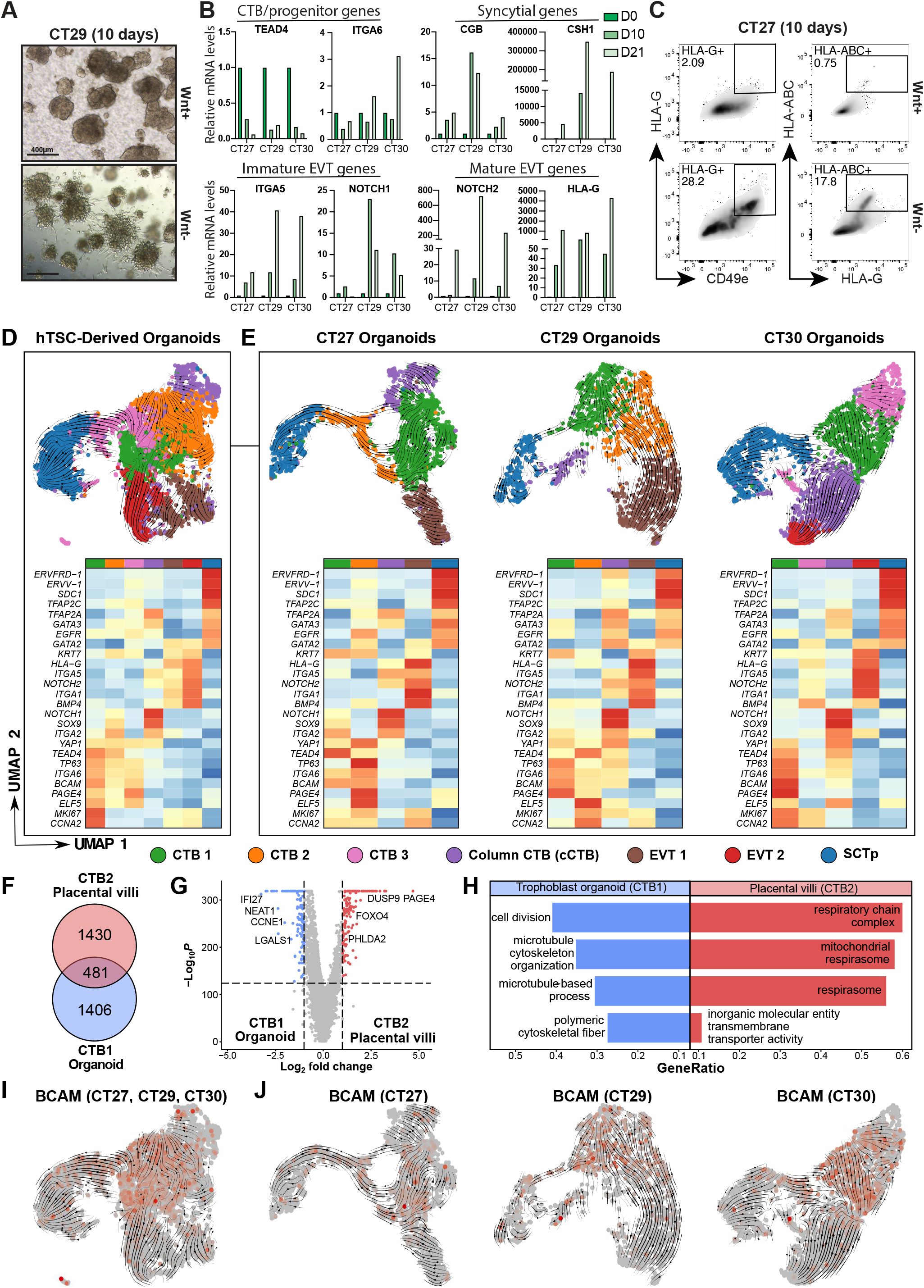
scRNA-seq analyses in hTSC organoids recapitulates a primitive CTB origin. **(A)** Bright field microscopy images of regenerative (Wnt+) and EVT-differentiated (Wnt-) hTSC-derived organoids following 10 days of culture. Bar = 400 μm. **(B)** Relative measurement of mRNA levels by qPCR in EVT-differentiated hTSC-derived organoids (CT27, CT29, CT30) over 21 days (D) of culture. Shown are CTB/progenitor (*TEAD4, ITGA6*), syncytial (*CGB, CSH1*), immature EVT (*ITGA5, NOTCH1*), and mature EVT (*NOTCH2, HLA-G*) genes. Each hTSC line was differentiated as one (n=1) replicate. **(C)** Representative flow cytometry plots showing cell frequencies of HLA-G, CD49e, and HLA-ABC positive cells in regenerative (Wnt+) and EVT-differentiated hTSC-derived (CT27) organoids. CT27 differentiation was performed on three (n=3) independent times. **(D)** UMAP and gene heatmap plots of combined CT27, CT29, and CT30 single cell hTSCs following culture in regenerative (Wnt+) and EVT-differentiated (Wnt-) conditions after 7 days in culture. UMAP plots are overlain with projected RNA velocity (scVelo) vector field, demonstrating 7 distinct trophoblast clusters representing 3 states (CTB1, CTB2, CTB3, bipotent cCTB, EVT1, EVT2, SCTp). Heatmap shows the average expression of trophoblast- and proliferation-associated genes in each cell state. **(E)** UMAP and gene heatmap plots derived from individual hTSC lines (CT27, 2259 cells; CT29, 1446 cells; CT30, 2604 cells) subjected to scRNA-seq analyses. UMAP plots are overlain with projected RNA velocity vector field, demonstrating 5 distinct trophoblast clusters in each hTSC line. **(F)** Venn diagram comparing genes shared and distinct between CTB2 in chorionic villi and CTB1 in hTSC organoids. **(G)** Volcano plot showing differentially expressed genes between CTB2 in chorionic villi and CTB1 in organoids. **(H)** Top pathways enriched in CTB2 (villi) and CTB1 (organoid) gene signatures. UMAP plots showing expression of BCAM (red) and velocity arrows in **(I)** combined (CT27, CT29, CT30) and **(J)** segregated hTSC-derived organoid cells/states.

Regenerative and EVT-differentiated hTSC organoid cells were then used to generate single cell cDNA libraries following 10X Genomics Chromium workflow. Combining single cell transcriptomic data from CT27, CT29, and CT30 organoids resolved 7 distinct cell states; three CTB (CTB1-3), a precursor to syncytiotrophoblast (SCTp), a putative column/EVT progenitor (cCTB), and two EVT-like states (EVT1, EVT2) (Fig. 3D). Hallmark gene markers of trophoblast-lineage (*TFAP2A, TFAP2C*), progenitor CTB (*TEAD4, YAP1, TP63, ITGA6, ELF5*), syncytiotrophoblast (*ERVFRD-1, ERVV-1, SDC1*), immature (*ITGA5, ITGA2, NOTCH1*) and mature EVT (*ITGA1, NOTCH2, HLA-G*), as well as gene markers of proliferation (*MKI67, CCNA2*) are shown corresponding to individual cell states (Fig. 3D; Table S2). CTB1-3 shared moderate overlap in gene expression, though CTB1 distinctly co-expressed proliferative and progenitor genes (Fig. 3D). While CTB2 and CTB3 also expressed progenitor genes, levels were comparatively lower than levels in CTB1 (Fig. 3D). The putative column/EVT progenitor state, cCTB, uniquely expressed *ITGA2, SOX9*, and the immature EVT marker *NOTCH1*; at the gene level, this unique state resembles the EVT progenitor population described by Lee *et al*. (2018) (Fig. 3D). States resembling terminally differentiated cell types showed strong enrichment for syncytiotrophoblast- and EVT-associated genes, with EVT1 showing the highest levels of *HLA-G* and *NOTCH2* (Fig. 3D).

When scRNA-seq data for hTSC organoids are plotted individually, overall cell state complexity reduces from 7 to 5 states (Fig. 3E). Intercomparison of hTSC lines showed notable variances in cell heterogeneity (Fig. 3E). While defined SCTp and EVT endpoint states are evident in all three lines, variances defined by gene expression in CTB and cCTB states suggest that the hTSC lines have distinct cell differentiation kinetics or pathways unique to themselves (Fig. 3E). Nonetheless, within each hTSC organoid, scVelo revealed a common CTB origin aligning to CTB1 (Fig. 3E). Similar to scVelo modeling in chorionic villi, 3D rendering of scVelo trajectories in organoids highlight multiple pathways extending from CTB1 to EVT, while a more direct single path is shown for SCTp establishment extending from CTB3 (Fig. S4). Examining gene signatures within these origins showed consistent enrichment of the trophoblast progenitor genes *TP63* and *YAP1*. However, we did see differences in levels of *ELF5* between hTSC lines. Specifically, *ELF5* was enriched in CTB2 in the CT27 and CT29 hTSC lines but aligned more with the origin and the CTB1 state in the CT30 hTSC line (Fig. 3E). Similar differences were observed for the proliferative gene markers *MKI67* and *CCNA2*, where levels were enriched within CTB1 of CT27 and CT30 but aligned more specifically to CTB2 in CT29 (Fig. 3E). When averaged across all hTSC lines, however, we observed consistent expression of *ELF5, BCAM, MKI67*, and *CNNA2* within the CTB1 state that overlapped with the predicted origin (Fig. 3D). Comparison of transcriptional signatures between the scVelo-determined origins in CTB from chorionic villi and in hTSC organoids highlights that there is considerable overlap between CTB2 (*in vivo*) and CTB1 (*in vitro*), as well as notable differences (Fig. 3F; Table S3). 481 genes are shared, whereas 1430 and 1406 genes are enriched within the origin states of chorionic villi and organoids, respectively (Fig 3F). A differential gene expression analysis between CTB2 (villi) and CTB1 (organoids) revealed 13,364 differentially expressed genes (Fig. 3G; Table S4). Pathway analysis of these signatures shows that organoid progenitors have enrichment of pathways underlying cell division and microtubule assembly, indicative of a pro-proliferative state, while villous progenitors showed enrichment of genes related to respiration, mitochondrial function, and inorganic transporter activity (Fig. 3H). Consistent with data from primitive CTB in chorionic villi, *BCAM* was shown to be one of the most highly expressed and FDR-ranked genes shared between organoid CTB1 and villous CTB2 states (Table S3), and also highly overlapped with the scVelo-predicted origin (Fig. 3I, 3J). These data show that molecular programs are largely conserved in hTSC organoids, that similarities in overall trophoblast heterogeneity are maintained, and that cell adhesion molecule *BCAM* is also a shared marker of regenerative CTB in chorionic villous and hTSC organoids.

### BCAM confers increased CTB regenerative potential

To our knowledge, BCAM localization within the human placenta has never been reported. To this end, we probed early (6.3 week) and late (11.2 week) first trimester placental villi specimens with BCAM antibody (Fig. 4A). A specific and strong BCAM signal was observed in CTB of both developmental time-points, and serial sections stained with keratin-7 (KRT7) and HLA-G showed that BCAM was present within proximal column cells, though the intensity of the signal was less than in villous CTB (Fig. 4A). Serial sections of chorionic villi stained with KRT7 and hCG clearly showed that BCAM is absent from the syncytiotrophoblast outer layer (Fig. 4A). Consistent with this, in hTSC-derived organoids cultured in regenerative conditions, BCAM signal was specific to the outer CTB layer and absent within the inner hCG^+^ syncytium (Fig. 4B). Dual labeling with BCAM and Ki67 antibody showed that BCAM^+^ CTB also co-express the proliferation marker (Fig. 4B).

**Figure 4:**
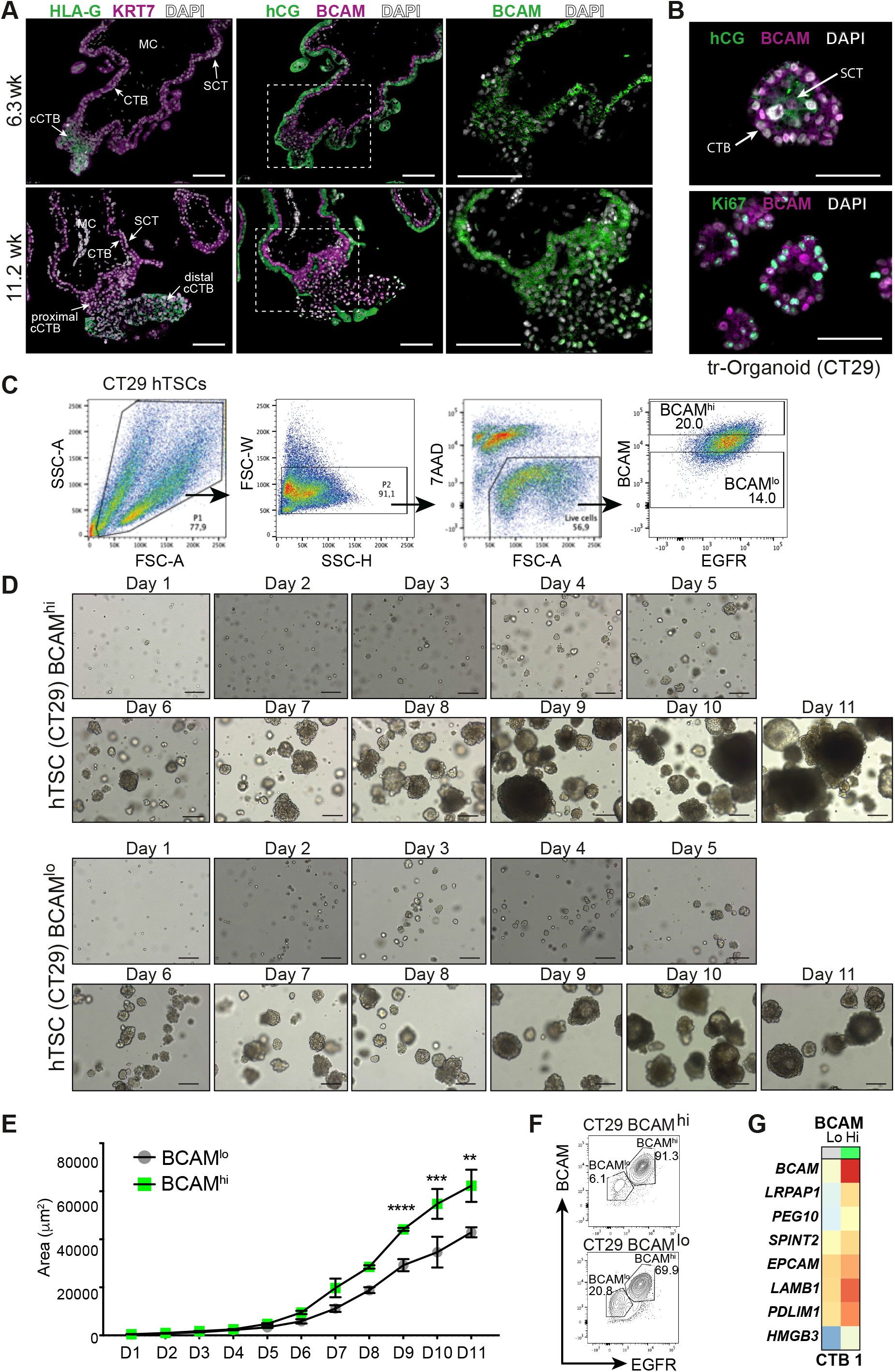
BCAM enrichment establishes a growth advantage in trophoblast organoids. **(A)** Immunofluorescence (IF) images of BCAM and keratin-7 (KRT7), human chorionic gonadotrophin (hCG), and HLA-G in early (6.3 weeks) and late (11.2 weeks) first trimester placental villi. Nuclei are stained with DAPI (white). Bars = 100μm. **(B)** IF images of BCAM, hCG and Ki67 in hTSC-derived organoid (CT29) cultured in regenerative (Wnt+) media for 7 days. **(C)** Cell sorting workflow for isolating BCAM^hi^ and BCAM^lo^ EGFR+ CTBs in hTSC (CT29)-derived organoids. **(D)** Bright field images of hTSC (CT29) BCAM^hi^ and BCAM^lo^ –derived organoids cultured over 11 days in regenerative Wnt+ media. Bars = 200μm. **(E)** Colony area (μm^2^) of BCAM^hi^ and BCAM^lo^ hTSC (CT29)-derived organoids cultured over 11 days (D). Each condition was performed on three (n=3) independent occasions. Median values are shown and statistical analyses were performed using a 2way ANOVA, multiple comparisons were controlled for using a Bonferroni post-test; significance *p* <0.05. ** = *p* < 0.01; **** = *p* <0.0001. **(F)** Flow cytometric panels showing expression of BCAM and EGFR in BCAM^hi^ and BCAM^lo^, CT29-derived organoids following 11 days of culture. **(G)** Heatmap showing levels of select differentially expressed genes from CTB expressing high (hi) or low (lo) levels of *BCAM* in the scRNA-seq defined CTB1 state of CT29-derived organoids following 7 days of culture.

To test the regenerative potential of BCAM-expressing cells, BCAM^hi^ and BCAM^lo^ CTBs were sorted from CT29 hTSC organoids, seeded at a density of 125 cells/μl within Matrigel domes, and cultured over 11 days (Fig. 4C, 4D). While the colony take-rate did not differ between BCAM^hi^ and BCAM^lo^ CTB organoids (Fig. S5), the colony size of BCAM^hi^ organoids appeared to be larger by day 7, with BCAM^hi^ -derived colonies being significantly larger than BCAM^lo^ -derived colonies on days 9-11 in culture (Fig. 4D, 4E). Flow cytometry assessment of day 11 BCAM^hi^ - and BCAM^lo^ -derived organoids showed that >90% of BCAM^hi^-sorted cells retained high levels of BCAM surface expression (Fig. 4F), while the frequency of CTBs expressing high levels of BCAM in BCAM^lo^-sorted cells was 70%, suggesting that the BCAM phenotype is dynamic. While a minor EGFR^lo^ population was detected in endpoint organoids from BCAM^hi^-sorted CTBs, a significant expansion in this population was evident in organoids derived from BCAM^lo^-sorted CTBs, suggesting that BCAM^lo^ CTB do not retain regenerative properties as robustly as BCAM^hi^ CTB (Fig. 4F). Comparing differentially expressed genes between CTB expressing high or low levels of *BCAM* in organoid scRNA-seq data showed that elevated *BCAM* levels associate with higher levels of the paternally imprinted transcription factor *PEG10*, and the stem cell renewal genes *EPCAM* and *HMGB3* (Nemeth, Kirby and Bodine, 2006) (Fig. 4G).

To specifically examine if BCAM plays a functional role in promoting trophoblast organoid growth, two *BCAM*-targeting siRNAs were transfected into CT29 hTSCs prior to and during organoid development, and organoid growth was monitored over 7 days. Non-transfected and non-silencing siRNA-transfected (siCTRL) CT29 hTSCs served as controls. siRNA-directed knockdown of BCAM showed a 40-60% reduction in BCAM protein levels at days 3 (mid) and 7 (endpoint) of culture (Fig. 5A, 5B). While non-transfected and siCTRL organoids formed large and robust colonies, BCAM silencing led to a consistent reduction in organoid colony size (Fig. 5C). Whereas control hTSCs formed greater frequencies of medium (2.0 - 4.9 ×10^4^ μm^2^) and large (> 5.0 ×10^4^ μm^2^) organoids, BCAM silenced hTSCs formed greater frequencies of small (< 2.0 ×10^4^ μm^2^) colonies (Fig. 5D). Similar to our observation that BCAM^lo^ sorted CTB showed an enhanced expansion of a EGFR^lo^ population, BCAM silencing in organoids led to a modest but significant increase in the frequency of EGFR^lo^ trophoblasts, suggesting that a reduction in BCAM may result in greater plasticity or accelerated differentiation (Fig. 5E). We also examined if BCAM impacts cell cycle kinetics, and by assessing DNA content using flow cytometry we show that BCAM silencing results in a modest, but not significant, increase in the proportion of trophoblasts in G_2_ phase, and a reciprocal, but not significant, decrease in the frequency of cells in S phase (Fig. 5F). Similarly in day 7 endpoint organoids, BCAM-silencing showed a reduction in the proportion of BrdU^+^ cells compared to frequencies in non-silencing controls, though this too did not reach statistical significance, likely attributed to variability in organoid growth and BrdU assessment of 3D cultures by IF (Fig. 5G).

**Figure 5:**
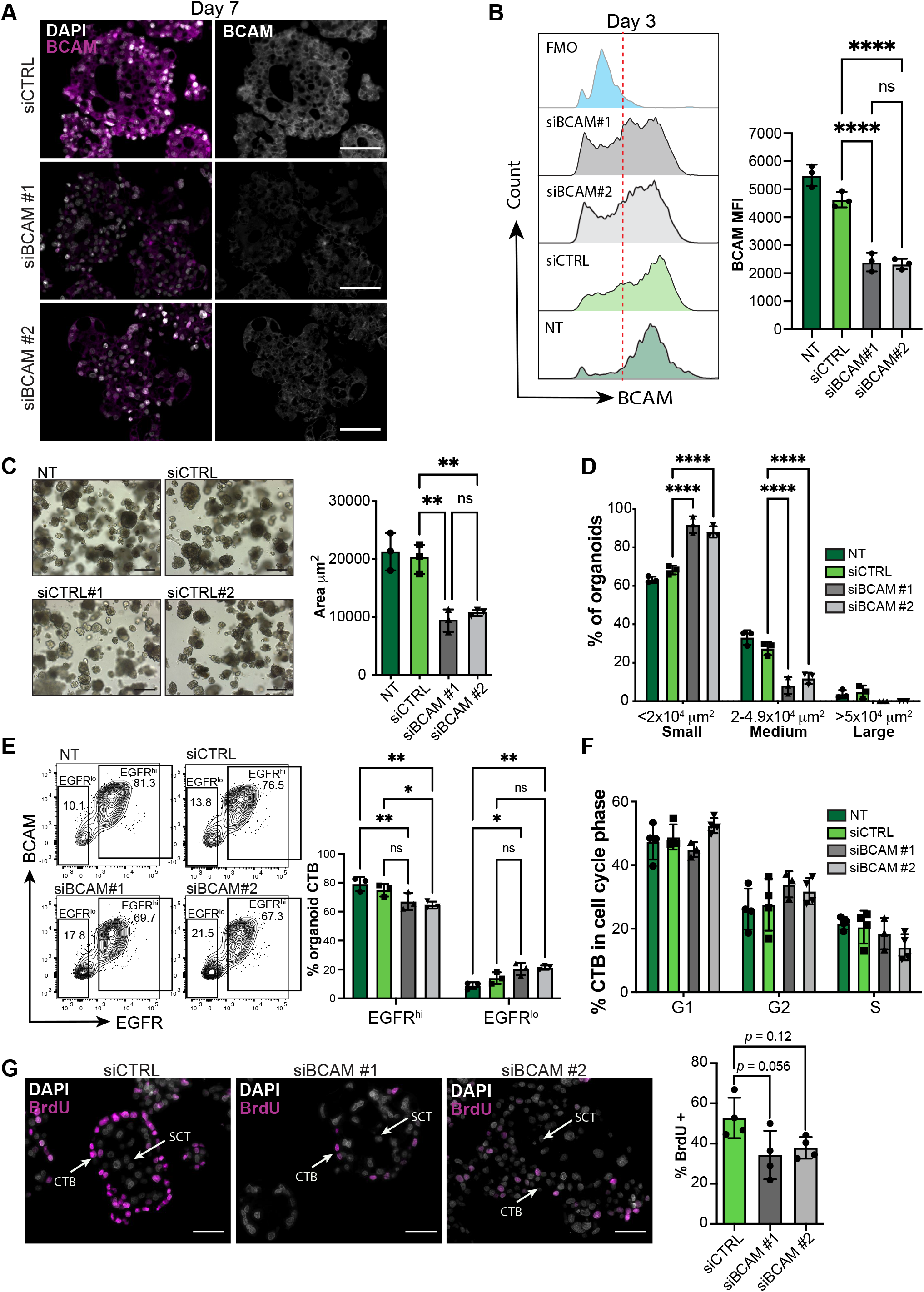
BCAM promotes CTB organoid growth. **(A)** IF images of BCAM in CT29 organoids transfected with control siRNA (ciCTRL) and BCAM-targeting siRNAs (siBCAM#1, siBCAM#2). Nuclei are stained with DAPI. Bars = 100 μm. **(B)** Representative flow cytometry plots showing BCAM levels in control (NT, siCTRL) and BCAM-silenced (siBCAM#1, siBCAM#2) CT29 organoids. FMO = fluorescence minus one. Bar graph to the right shows quantification of BCAM median fluorescence intensity (MFI). Statistical analyses between groups were performed using ANOVA and two-tailed Tukey post-test; significance *p* <0.05. Each condition was performed on four (n=4) independent occasions. **(C)** Bright field images of CT29 organoids transfected with control (NT, siCTRL) or BCAM-silencing siRNA (siBCAM#1, siBCAM#2) following 7 days of culture in regenerative Wnt+ media. Bars = 200μm. Bar graph to the right shows area quantification of control and BCAM-silenced organoids. Median values are shown and statistical analyses between groups were performed using ANOVA and two-tailed Tukey post-test; significance *p* <0.05. Each condition was performed on three (n=3) independent occasions. **(D)** Bar graph shows proportions of small, medium, and large colonies in control- and BCAM-silenced CT29 organoids. Median values are shown and statistical analyses were performed using a 2way ANOVA, multiple comparisons were controlled for using a Bonferroni post-test; significance *p* <0.05. **(E)** Flow cytometry plots showing EGFR and BCAM frequencies in control and BCAM-silenced CT27 organoids at day 3 of organoid culture in regenerative media. Bar graphs show median values of EGFR^hi^ and EGFR^lo^ frequencies, and statistical analyses between groups were performed using ANOVA and two-tailed Tukey post-test; significance *p* <0.05. Each condition was performed on three (n=3) independent occasions. **(F)** Bar graphs show frequency of cell G_1_, G_2_, and S cell cycle phases measured by flow cytometry in control (NT, siCTRL) and BCAM-silenced (siBCAM#1, siBCAM#2) CT29 organoids. Statistical analyses between groups were performed using ANOVA and two-tailed Tukey post-test; significance *p* <0.05. **(G)** IF images showing Brdu incorporation in control (siCTRL) or BCAM-silenced day 7 organoids following 4 hr of BrdU pulse. Bar graphs show median values of BrdU frequencies in each group. Bars = 200μm. Each condition was performed on three or four (n=3, n=4) independent occasions. * = *p* < 0.05; ** = *p* < 0.01; **** = *p* <0.0001.

## DISCUSSION

Here we applied single cell RNA sequencing of human placentas at two gestational age ranges within the first trimester to further our understanding of human trophoblast development. In doing this, we provide here novel transcriptomic data from 7,798 human trophoblasts, resolve the first trimester trophoblast landscape at increased resolution, and clarify how trophoblast heterogeneity changes during the first trimester. Using state-of-the-art single cell lineage trajectory tools, we trace back the lineage commitments made by developing CTBs to identify a trophoblast progenitor origin marked by increased expression of the cell surface glycoprotein BCAM. BCAM’s association with the predicted CTB origin highlights an additional utility of using BCAM as a new cell surface progenitor marker for future studies identifying and defining CTB progenitors.

Previous studies have explored the placental transcriptome at single cell resolution (Liu *et al*., 2018; Suryawanshi *et al*., 2018; Vento-Tormo *et al*., 2018). While these studies have contributed important insight into placental cell heterogeneity, a strict focus on understanding CTB hierarchy and gene pathways underlying CTB regeneration and commitment to EVT or SCT was not a focus of these studies. Our study identifies four unique CTB states that are distinguishable by state of metabolism (protein translation, CTB1; mitochondrial regulation, CTB3), cell cycle (CTB4), and molecular readouts of respiration and transporter activity (CTB2). Concurrent single cell analyses using hTSC-derived organoids identifies similar cell states found in vivo, though differences in the number of cell state clusters and specific genes in organoids derived from unique hTSC lines does suggest CTB clusters may represent snap-shots of transient cell states captured during CTB maintenance and differentiation, and are not specific cell types *per se*.

Using scVelo and Monocle3, we identified a CTB origin that likely represents a primitive stem cell-like niche. CTB2 was identified by both scVelo and Monocle3 trajectory modeling tools as the CTB state overlapping with this predicted origin. We were initially surprised to see the predicted origin did not directly align with CTB3, as this state showed robust levels of many classical CTB progenitor genes like *YAP1, TEAD4*, and *TP63*, where by comparison CTB2 expressed lower levels of these known progenitor genes. However, CTB2 did show enrichment for some important CTB-associated genes like *SPINT1* (Walentin *et al*., 2015), and further showed elevated levels for the Wnt ligand *WNT7A* and other Wnt-modifying factors like *GPC3, DAB2*, and *DACT2* (Jiang, He and Howe, 2012; Capurro *et al*., 2014; Wang *et al*., 2015), suggesting that this state is responsive to WNT signaling, an important feature underlying *in vitro* CTB progenitor maintenance (Okae *et al*., 2018).

Comparisons between our placental tissue sequencing data and our trophoblast organoid sequencing data demonstrate similarities in general CTB, SCT and EVT function, but also subtle differences between the different states of CTB found in each stem cell derived organoid condition and our tissue CTBs. Comparison of the CTB progenitor niche identified in both the organoid and tissue datasets identifies the common transcription factors *YAP1, SPINT2, PEG10*, and *MSI1* as well as consistent upregulation of *BCAM*, supporting identification of BCAM as a CTB progenitor marker. Interestingly, quantification of mRNA levels of known CTB related genes (*TEAD4* and *ITGA6*) as well as SCT (*CGB* and *CSH1*) and EVT related genes (*ITGA5, NOTCH1, NOTCH2*, and *HLA-G*) on days 0, 11, and 21 of differentiation indicate that hTSC specific organoid colonies show different rates of differentiation or capacities to differentiate. Our EVT-differentiated organoid transcriptomes were sequenced on day 7 of differentiation, indicating that the differences we are observing may in part be due to groups of organoid colonies being at a different stages of differentiation, with CT29 derived organoids showing the most rapid capacity for EVT differentiation, followed by CT30 and CT27. These differences in differentiation kinetics or capacities should be taken into consideration by researchers aiming to use these regenerative trophoblast lines for future research.

Consistent between chorionic villous and organoid datasets is the identification of two CTB progenitor states in the first trimester placenta, one representing a CTB origin state marked by elevated *BCAM*, and one representing a more specified bipotent EVT progenitor cell state, marked by *ITGA2* and *NOTCH1*. In 8-9 week GA placental villi, ITGA2 is specific to proximal cells within anchoring placental villi near the basement membrane (Lee *et al*., 2018). It is thought that as chorionic villi develop, trophoblast progenitors within close proximity to uterine stroma adopt a unique gene expression profile that predisposes their differentiation along the extravillous pathway. We show that the transcriptome of *ITGA2*^*+*^ putative column progenitors express EVT-associated genes such as *NOTCH1* and *ITGA5*, as well as the CTB/progenitor-associated genes *GATA3* and *ELF5*. ITGA2^+^ cCTBs are described as having a level of bipotency. Analyses using scVelo suggests *ITGA2*^+^column/EVT progenitors descend from the *BCAM*^+^ CTB progenitor origin and are maintained as a reserve progenitor population, though whether this state represents a bipotent CTB state will need to be fully examined, but nonetheless our analyses demonstrate a detailed molecular signature for this interesting EVT progenitor state.

We identify *BCAM* as a gene expressed by CTB progenitors, where its expression is particularly elevated in the most upstream CTB states. BCAM interacts predominantly with its ligand *LAMA5*, a component of the laminin-511 heterotrimers found within the placental extracellular matrix (Zen *et al*., 1999; Kikkawa *et al*., 2013). BCAM is an antigen upregulated in ovarian carcinoma, and facilitates tumor migration by competing for laminin binding with integrins (Kannan *et al*., 2015; Guadall *et al*., 2019). Here, we show that flow-cytometry enrichment of BCAM in CTBs significantly potentiates CTB organoid growth. Interestingly, no differences were observed in the capacity of BCAM^hi^ or BCAM^lo^ CTBs to form organoid colonies from single cells, suggesting that BCAM^hi^ CTB may not have intrinsically greater stem-like properties over BCAM^lo^ CTB. Rather, BCAM may enhance organoid growth by potentiating survival signals induced by cell-ECM engagement, or by initiating/driving CTB proliferation pathways. However, we show that genes associated with stem cell regeneration are elevated in BCAM^hi^ hTSC-derived organoids as compared to organoids derived from BCAM^lo^ cells. Specifically, we show increased expression of *PEG10, EPCAM*, and *HMGB3*, indicating that BCAM may confer increased organoid growth by controlling stem cell maintenance or regeneration. In our experimental organoid system, even though BCAM^lo^ organoids form smaller colonies by day 11, assessment of BCAM expression in BCAM^lo^ organoids at endpoint shows that BCAM levels and the frequency of BCAM-expressing cells is comparable to BCAM^hi^ CTBs. This suggests that BCAM expression in progenitor CTBs is dynamic, and that requirements for growth, stemness, or survival result in the induction of BCAM expression. Consistent for a role for BCAM in controlling organoid growth, BCAM silencing also resulted in the formation of smaller organoids, where cell cycle analysis suggests that BCAM regulates progenitor entry/exit into the cell cycle. Future research should focus on how BCAM potentially creates a supportive stem/regenerative cell niche, possibly by mediating specific cell-ECM engagement signals important in progenitor maintenance.

The identification of a novel cell surface human CTB progenitor cell marker provides improved methods for isolating and studying the mechanisms regulating human CTB progenitor cells, trophoblast differentiation, and human placental development. We have used single cell RNA sequencing to define trophoblast differentiation trajectories in early human placental development. Importantly, we identify BCAM as a new CTB progenitor marker that confers enhanced clonal growth. Future work is needed to clarify how increased BCAM expression facilitates improved CTB growth, and the exact molecular paths through which this occurs. We hope that future experiments will be able to capitalize on the identification of BCAM as a cell surface CTB progenitor marker through which these cells can be isolated and further studied.

## MATERIALS AND METHODS

### Patient recruitment and tissue collection

Placental tissues were obtained with approval from the Research Ethics Board on the use of human subjects, University of British Columbia (H13-00640). All samples were collected from women (19 to 39 years of age) providing written informed consent undergoing elective terminations of pregnancy at British Columbia’s Women’s Hospital, Vancouver, Canada. First trimester placental tissues (*n*=9) were collected from participating women (gestational ages ranging from 5–12 weeks) having confirmed viable pregnancies by ultrasound-measured fetal heartbeat. Patient clinical characteristics i.e. height and weight were additionally obtained to calculate body mass index (BMI: kg/m^2^) and all consenting women provided self-reported information via questionnaire to having first hand exposure to cigarette smoke or taken anti-inflammation or anti-hypertensive medications during pregnancy. Patient and sample phenotype metadata are summarized in Supplemental Table 1.

### Human trophoblast organoids

#### Establishment and culture

Human trophoblast stem cells (hTSC) CT27 (RCB4936), CT29 (RCB4937), and CT30 (RCB4938) from Riken BRC Cell Bank, were established from first trimester CTBs by Okae et al (Okae *et al*., 2018) and used here to generate organoid cultures using a modification of previously described protocols (Haider *et al*., 2018; Turco *et al*., 2018). In summary, these cells were re-suspended in ice-cold growth factor-reduced Matrigel (GFR-M, Corning) to a final concentration of 100%. Cell suspensions (40 µl) containing 5-20 × 10^3^ cells were carefully seeded in the center of each well of a 24-well plate. Plates were placed at 37°C for 2 minutes. Following this, the plates were turned upside down to promote equal spreading of the cells throughout the Matrigel domes. After 15 minutes, the plates were flipped back to normal and the Matrigel domes were overlaid with 0.5 mL pre-warmed trophoblast organoid media containing advanced DMEM/F12 (Gibco), 10 mM HEPES, 1 x B27 (Gibco), 1 x N2 (Gibco), 2 mM L-glutamine (Gibco), 100 µg/ml Primocin (Invivogen), 100 ng/mL R-spondin (PeproTech), 1 µM A83-01 (Tocris), 100 ng/mL rhEGF (PeproTech), 3 µM CHIR99021 (Tocris), and 2 µM Y-27632 (EMD Millipore). Organoids were cultured for 7-21 days.

#### EVT Differentiation

EVT differentiation of trophoblast organoids was performed using an adapted of previously published protocols (Okae *et al*., 2018). In summary, organoids were cultured until > 50% colonies reached at least 100 µm in diameter (∼3-5 days). Following this, organoids were maintained in EVT differentiation media containing advanced DMEM/F12, 0.1 mM 2-mercaptoethanol (Gibco), 0.5% penicillin/streptomycin (Gibco), 0.3% BSA (Sigma Aldrich), 1% ITS-X supplement (Gibco), 100 ng/mL NRG1 (Cell Signaling Technology), 7.5 µM A83-01 (Tocris), 4% Knockout Serum Replacement (Gibco). Media exchanged for EVT differentiation media lacking NRG1 around day 7-10 of differentiation following the visible outgrowth of cells. Differentiation was stopped at day 7 (D7) for 10X single cell RNA sequencing and flow cytometry analysis. Differentiation was stopped at day 21 (D21) for qPCR analysis.

### Placental tissue and trophoblast organoid single cell RNA-seq library construction

#### Placental tissue

Healthy placental villi were physically scraped form chorionic membrane by scalpel and enzymatically digested in 0.5 ml collagenase/hyaluronidase and 1.5 ml DMEM/F12 with 500 μl penicillin/streptomycin and 500 μl 100X antibiotic/antimycotic at 37°C for 45 minutes. Placental villi cell suspensions were diluted in 1X Hank’s Balanced Salt Solution (HBSS) containing 2% FBS, pelleted at 1200 rpm for 5 minutes, and resuspended in 0.25% trypsin-EDTA. The solution was diluted again in 1X HBSS containing 2% FBS, pelleted at 1200 rpm for 5 minutes, and resuspended in dispase and 1 mg/ml DNase I. Liberated cells were enriched by passing digest through through a 40 μm filter.

#### Trophoblast organoids

For all organoid cultures, media was aspirated, and organoids were diluted in Cell Recovery Solution at 4°C for 40 minutes on either culture day 0 in complete media (non-differentiated organoids) or culture day 7 in differentiation media (EVT-differentiated organoids). Cells were then washed with cold PBS and centrifuged at 1000 rpm for 5 minutes. Cells were disaggregated in TrypLE containing 5% DNase I at 37°C for 7 minutes before the addition of 100 μl complete media. The resulting cell suspension was then sieved through a 40 μm filter.

#### Library construction

Single cell suspensions (tissue- and organoid-derived) were stained with 7-Aminoactinomycin D (7-AAD) viability marker and live/dead sorted using a Becton Dickinson (BD) FACS Aria. Cell suspensions were loaded onto the 10X Genomics single cell controller for library preparation. Placental villi-derived single cell suspensions were prepared using the chromium single cell 3’ reagent v2 chemistry kit, following standard manufacturer protocol for all steps. Non-differentiated and differentiated trophoblast organoid-derived single cell suspensions were prepared using the chromium single cell 3’ reagent v3 chemistry kit, following standard manufacturer protocol for all steps. Single cell libraries were sequenced on an Illumina Nextseq 500 instrument at a sequencing depth of 50,000 reads/cell for tissue-derived cells and 150,000 reads/cell for organoid-derived cells. For placental villus samples, the 10X Genomics Cell Ranger version 2.0 software was used for STAR read alignment and counting against the provided GRCh38 reference while the trophoblast organoid samples used the 10X Genomics Cell Ranger version 3.0 software for STAR read alignment and counting against the GRCh38 reference.

#### Data Repository Integration

Droplet-based first trimester scRNA-seq decidual and placental tissue data was obtained from the public repository ArrayExpress (available at: E-MTAB-6701) (Vento-Tormo *et al*., 2018). Raw BAM files were downloaded and pre-processed using 10X Genomics Cell Ranger version 2.0 under identical parameters as previously described for our placental villi and trophoblast organoid samples using the GRCh38 reference. Sample specific metrics are summarized in Supplemental Table 2.

### Single cell RNA-seq data analysis

#### Data pre-processing and quality control

A total of 50,790 single cells were sequenced and pre-processed using the Seurat R package (Butler *et al*., 2018; Stuart *et al*., 2019). To ensure quality cells were used in downstream analyses, cells containing fewer than 1000 detected genes and greater than 20% mitochondrial DNA content were removed. Individual samples were scored based on expression of G2/M and S phase cell cycle gene markers, scaled, normalized, and contaminating cell doublets were removed using the DoubletFinder package (McGinnis, Murrow and Gartner, 2019) with default parameters and a doublet formation rate estimated from the number of captured cells. Following pre-processing, all samples were merged and integrated using cell pairwise correspondences between single cells across sequencing batches. During integration, the Pearson residuals obtained from the default regularized negative binomial model were re-computed and scaled to remove latent variation in percent mitochondrial DNA content as well as latent variation resulting from the difference between the G2M and S phase cell cycle scores.

#### Cell clustering, identification, and trophoblast sub-setting

Cell clustering was accomplished in Seurat at a resolution of 0.90 using 50 principal components. Cell identity was determined through observation of known gene markers found within the maternal fetal interface (vimentin, CD45, neural cell adhesion molecule, CD14, etc.) and identifying cluster marker genes via the “FindAllMarkers” function using a model-based analysis of single-cell transcriptomics (MAST) GLM-framework implemented in Seurat (Finak *et al*., 2015). Gene markers were selected as genes with a minimum log fold change value > 0.20 and an adjusted p-value < 0.05 within a specific cluster. Trophoblasts were identified as expressing *KRT7, EGFR, TEAD4, TP63, TFAP2A, TFAP2C, GATA2, GATA3, HLA-G, ERVFRD-1* transcripts at a level greater than zero, and *VIM, WARS, PTPRC, DCN, CD34, CD14, CD86, CD163, NKG7, KLRD1, HLA-DPA1, HLA-DPB1, HLA-DRA, HLA-DRB1, HLA-DRB5, HLA-DQA1* transcripts at a level equal to zero. Trophoblasts were subset and re-clustered in Seurat at a resolution of 0.375 using 30 principal components. As mentioned previously, trophoblast cluster identity was determined through observation of known trophoblast gene marker expression, proliferative gene marker expression, and from unique cluster genes identified by differential expression analysis in Seurat. Trophoblast projections were visualized using Uniform Manifold Approximation and Projections (UMAPs).

#### Differential gene expression

Differential expression (DEG) analyses were performed in Seurat using the “FindMarkers” function using a MAST framework on the raw count data. Parameters were limited to genes with a log fold-change difference between groups of 0.20 and genes detected in a minimum 10% cells in each group. Results were filtered for positive gene markers and significant DEGs were taken as having a fold change difference > 0.25 and an adjusted p value < 0.05.

#### Gene set enrichment analysis

GSEA analyses were performed in Seurat using the “FindMarkers” function using a MAST framework on the raw count data. Parameters included a minimum log fold-change difference between groups of “-INF”, a minimum gene detection of “-INF” in cells in each group, and a minimum difference in the fraction of detected genes within each group of “-INF”, allowing all genes to be compared. Results were visualized using the R package EnhancedVolcano (version 1.8.0), with a fold-change cut-off of 0.25 and a p value cut off of 0.00005 taken as the thresholds for significance.

#### Gene ontology

GO analysis of significant differentially expressed genes was performed using the clusterProfiler R package (Yu *et al*., 2012) and the top enriched terms (Benjamini-Hochberg corrected p value <0.05) were visualized as dot plots using the R package enrich plot (version 1.10.2).

#### Pseudotime

The Monocle2 R package (Trapnell *et al*., 2014; Qiu, Hill, *et al*., 2017; Qiu, Mao, *et al*., 2017) was used to explore the differentiation of progenitor CTBs into specialized SCT and EVT. The trophoblast cell raw counts were loaded into Monocle2 for semi-supervised single cell ordering in pseudotime, taking increased *BCAM* expression to represent cells at the origin and *HLA-G* and *ERVFRD-1* gene expression to indicate progress towards EVT and SCT, respectively. Branched expression analysis modelling (BEAM) was applied to observe the expression of identified trophoblast cluster gene markers and genes related to the WNT, HIPPO, NOTCH, and EGFR signaling pathways over pseudotime, progressing towards either EVT or SCT. Results were visualized using the “plot_cell_trajectory” function. To resolve cytotrophoblast differentiation trajectories at higher resolution, the Monocle3 package (Cao *et al*., 2019) was utilized. Trophoblast raw counts were input, clustered, and the principal graph was created using the “learn_graph” function. The “order_cells” function was then used to order cells in pseudotime, designating the trophoblast root within the CTB 2 state.

#### Lineage Trajectory

The Velocyto package (La Manno *et al*., 2018) was run in terminal using the command line interface to generate sample-specific files containing count matrices of unspliced, spliced, and ambiguous RNA transcript abundances. The velocyto.R implementation of the package was then used to merge each file with its corresponding 10X sample during data pre-processing in Seurat. The final, processed trophoblast object and UMAP embeddings were exported to a Jupyter Notebook and the scVelo Python package (Bergen *et al*., 2020) was used to read, prepare, and recover holistic trophoblast gene splicing kinetics. Velocities were projected on top of the extracted UMAP embeddings generated in Seurat.

### Flow cytometry analysis and cell sorting

#### Organoid single cell workflow

Single cells from CT29 organoids were isolated by incubation with Cell Recovery Solution (Corning) for 40 minutes at 4°C. Then, cells were washed in ice-cold PBS and dissociated into single cells using TrypLE™ supplemented with 5% DNAse I for 7 minutes at 37°C.

#### Phenotyping EVT differentiated CT27, CT29, and CT30 organoids

Following single cell dissociation, organoid cells were stained with EGFR BV605 (clone AY13; 1:50; Biolegend), HLA-G APC (clone MEMG-9; 1:50; Abcam), HLA-ABC Pacific Blue (clone W6/32; 1:100; Biolegend), CD49e/ITGA5 PeCy7 (clone eBioGoH3; 1:200; Thermo Fisher Scientific), CD49f/ITGA6 Alexa 488 (clone P1D6; 1:200; Thermo Fisher Scientific), and BCAM BV711 (clone B64; 1:50; BD Biosciences). *Phenotyping BCAM*^*hi*^ *and BCAM*^*lo*^ *organoids*: Single cells were incubated with anti-human BCAM BV421 (clone B64; 1:50; BD Biosciences), EGFR BV605, and CD49f/ITGA6 PeCy7 for 30 minutes in the dark on ice, followed by viability staining with 7-AAD. Samples were analyzed using LSRII (Becton Dickinson) and a minimum of 20,000 events were acquired.

#### hTSC BCAM^hi/lo^ sorting

CT29 hTSCs were dissociated using TrypLE™ (Thermo Fisher Scientific) with 5% DNAse I (Sigma Aldrich) for 7 minutes at 37°C. Following resuspension in PBS supplemented with Antibiotic-Antimycotic (Gibco) and 1% P/S (Sigma Aldrich), cells were stained with BCAM BV421 (clone B64; 1:50; BD Biosciences) and EGFR Pecy7 (clone AY13; 1:50; Biolegend) for 30 minutes at 4°C in the dark, followed by incubation with 7-AAD (Thermo Fisher) for viability assessment. CT29 BCAM^hi^ and BCAM^lo^ populations were sorted using a FACS Aria (BD Biosciences).

#### Cell cycle analysis

Cell cycle phases were determined on day 3 of organoid culture using 7AAD assessment of DNA quantity. A minimum of 5,000 events were collected and cell cycle was analysed using Watson model on FlowJo 10.7.1.

### Trophoblast organoid growth assay

CT29 sorted BCAM^hi^ and BCAM^lo^ cells were seeded at a density of 5,000 cells in 40 µL of Matrigel in the presence of trophoblast organoid media. Phase contrast images were taken with AxioObserver inverted microscope (Carl Zeiss) using 20X objective, and organoids were further analyzed by qPCR, immunofluorescence, and flow cytometry. Organoid formation efficiency was assessed by measuring organoid number and size (area in μm^2^). Images were taken daily and analyzed using imageJ (NIH). The scale was set by measuring the scale bar in pixels (285.5 pixels = 200 μm).

### RNA isolation

Prior to RNA extraction, organoids were dissociated into single cells as described above. Total RNA was prepared from these cells using RiboZol reagent (VWR Life Science) followed by RNeasy Mini kit (Qiagen) according to manufacturer’s protocol. RNA purity was confirmed using a NanoDrop Spectrophotometer (Thermo Fisher Scientific).

### cDNA synthesis and qPCR

RNA was reverse transcribed using qScript cDNA SuperMix synthesis kit (Quanta Biosciences) and subjected to qPCR (ΔΔCT) analysis, using PowerUP SYBR Green Master Mix (Thermo Fisher Scientific) on a StepOnePlus Real-Time PCR System (Thermo Fisher Scientific) and forward and reverse primer sets for *TEAD4* (F: 5’-CGGCACCATTACCTCCAACG-3’; R: 5’-CTGCGTCATTGTCGATGGGC-3’), *ITGA6* (F: 5’-CACTCGGGAGGACAACGTGA-3’; R: 5’-ACAGCCCTCCCGTTCTGTTG-3’), *CGB* (F: 5’-GCCTCATCCTTGGCGCTAGA-3’; R: 5’-TATACCTCGGGGTTGTGGGG-3’), *CSH1* (F: 5’-GACTGGGCAGATCCTCAAGCA-3’; R: 5’-CCATGCGCAGGAATGTCTCG-3’), *ITGA5* (F: 5’-GGGTCGGGGGCTTCAACTTA-3’; R: 5’-ACACTGACCCCGTCTGTTCC-3’), *HLA-G* (F: 5’-CCCACGCACAGACTGACAGA-3’; R: 5’-AGGTCGCAGCCAATCATCCA-3’), *NOTCH1* (F: 5’-GCCTGAATGGCGGGAAGTGT-3’; R: 5’-TAGTCTGCCACGCCTCTGC-3’), *NOTCH2* (F: 5’-ACTCGGGGCCTACTCTGTGA-3’; R: 5’-TGTCTCCCTCACAACGCTCC-3’), *GAPDH* (F: 5’-AGGGCTGCTTTTAACTCTGGT-3’ R: 5’-CCCCACTTGATTTTGGAGGGA-3’). All raw data were analyzed using StepOne Software v2.3 (Thermo Fisher Scientific). The threshold cycle values were used to calculate relative gene expression levels. Values were normalized to endogenous *GAPDH* transcripts.

### Immunofluorescence microscopy

Placental villi (6-12 weeks’ gestation; n=2) were fixed in 2% paraformaldehyde overnight at 4 ° C. Tissues were paraffin embedded and sectioned at 6 μm onto glass slides. Immunofluorescence was performed as previously described. Briefly, organoids or placental tissues underwent antigen retrieval by heating slides in a microwave for 5 × 2-minute intervals in citrate buffer (pH 6.0). Sections were incubated with sodium borohydride for 5 minutes at room temperature (RT), followed by Triton X-100 permeabilization for (Aghababaei *et al*., 2015) 5 minutes, RT. Slides were blocked in 5% normal goat serum/0.1% saponin for 1hr, RT, and incubated with combinations of the indicated antibodies overnight at 4 °C: mouse monoclonal anti-HLA-G (clone 4H84; 1:100; Exbio, Vestec, Czech Republic); rabbit monoclonal anti-cytokeratin 7 (clone SP52; 1:50; Ventana Medical Systems), rabbit polyclonal anti-BCAM (1:100; Abcam); mouse monoclonal anti-hCG beta (clone 5H4-E2; 1:100; Abcam), anti-BrdU (1:1000, Bu20a, Cell Signaling Technology). Following overnight incubation, sections and coverslips were washed with PBS and incubated with Alexa Fluor goat anti-rabbit 568 and goat anti-mouse 488 conjugated secondary antibodies (Life Technologies, Carlsbad, CA) for 1h at RT, washed in PBS and mounted using ProLong Gold mounting media containing DAPI (Life Technologies).

Slides were imaged with an AxioObserver inverted microscope (Car Zeiss, Jena, Germany) using 20X Plan-Apochromat/0.80NA or 40X Plan-Apochromat oil/1.4NA objectives (Carl Zeiss). An ApoTome .2 structured illumination device (Carl Zeiss) set at full Z-stack mode and 5 phase images was used for image acquisition.

### siRNA knockdown of *BCAM*

Two BCAM-targeting siRNAs (siBCAM #1, siBCAM #2; Dharmacon) were transfected into CT29 hTSCs using Lipofectamine RNAiMAX (Thermo Fisher) at a concentration of 100 nM. For controls, cells were transfected with ON-TARGETplus non-silencing siRNA (siCTRL; Cat# D-001810-01-20; Dharmacon) or cultured in the presence of transfection reagent alone (non-treated; NT). CT29 hTSCs cultured in 2D were initially forward-transfected using Lipofectamine RNAiMAX (Thermo Fisher) to achieve efficient *BCAM* silencing prior to organoid establishment. Following 48 h, hTSCs cells were transfected with BCAM and CTRL siRNAs for a second time following a reverse transfection approach and embedded in Matrigel to initiate 3D trophoblast organoids. Following 3- or 7-days post-transfection, organoids were either cell dissociated from Matrigel using Cell Recovery Solution (BD Biosciences) for BCAM cell surface quantification using flow cytometry, or fixed in 4% paraformaldehyde, embedded in paraffin, and sectioned onto glass slides for IF assessment of BCAM, respectively.

### BrdU incorporation assay

Pulse-chase labeling with BrdU (Sigma-Aldrich) was conducted on CT29 hTSC-derived organoids. Two days following 3D hTSC organoid establishment and reverse control (siCTLR) and BCAM siRNA (siBCAM#1, siBCAM#2) transfection, organoids were exposed to a 4-hr pulse with culture media containing 10 μM of BrdU. Following 4 hr of labelling, organoids were washed in PBS and fixed in 4% PFA overnight. Organoids were paraffin embedded and sectioned for immunofluorescence microscopy. Immunofluorescent staining with anti-BrdU antibody (1:200, Bu20a, Sigma-Aldrich) with the addition of a 30-minute incubation in 2M hydrochloric acid between permeabilization and sodium borohydride steps.

### Statistical analysis

Data are reported as median values with standard deviations. All calculations were carried out using GraphPad Prism 9 software (San Diego, CA). For single comparisons, Mann-Whitney non-parametric unpaired t-tests were performed. For multiple comparisons, one-way Kruskal-Wallis ANOVA followed by Dunn’s multiple comparison test was performed. One-way ANOVA followed by Tukey post-test were performed for all other multiple comparisons. The differences were accepted as significant at *P* < 0.05. For scRNA-seq statistical analyses, please refer to scRNA-seq analysis sections in methods.

## DATA AVAILABILITY/ACCESSION NUMBER

The ArrayExpress accession number for the publicly available data reported in this paper is: E-MTAB-6701

## AUTHOR CONTRIBUTIONS

AGB designed the research. AGB, MS, JB, BC, JW, SY, JT, HTL, PML performed experiments and analysed data. AGB, MS, JB, and BC wrote the paper. All authors read and approved the manuscript.

## ACKNOWLEDGEMENTS

The authors extend their sincere gratitude to the hard work of staff at British Columbia’s Women’s Hospital’s CARE Program for recruiting participants to our study and the staff at the University of British Columbia Biomedical Research Centre, especially Yiwei Zhao and Ryan Vander Werff, for assistance with all drop-seq library preparation and sequencing.

## Abbreviations

CTB: Cytotrophoblast
DCT: Distal column trophoblast
DGE: Differential gene expression
EVT: Extravillous trophoblast
FDR: False discovery rate
GO: Gene ontology
HLA-G: Human leukocyte antigen G
Hr: Hour
RT: Room Temperature
IF: Immunofluorescence
scRNA-seq: single cell RNA sequencing
SCT: Syncytiotrophoblast
UMAP: Uniform manifold approximation and projection

## TITLES AND LEGENDS TO FIGURES

**Supplemental Figure 1:**
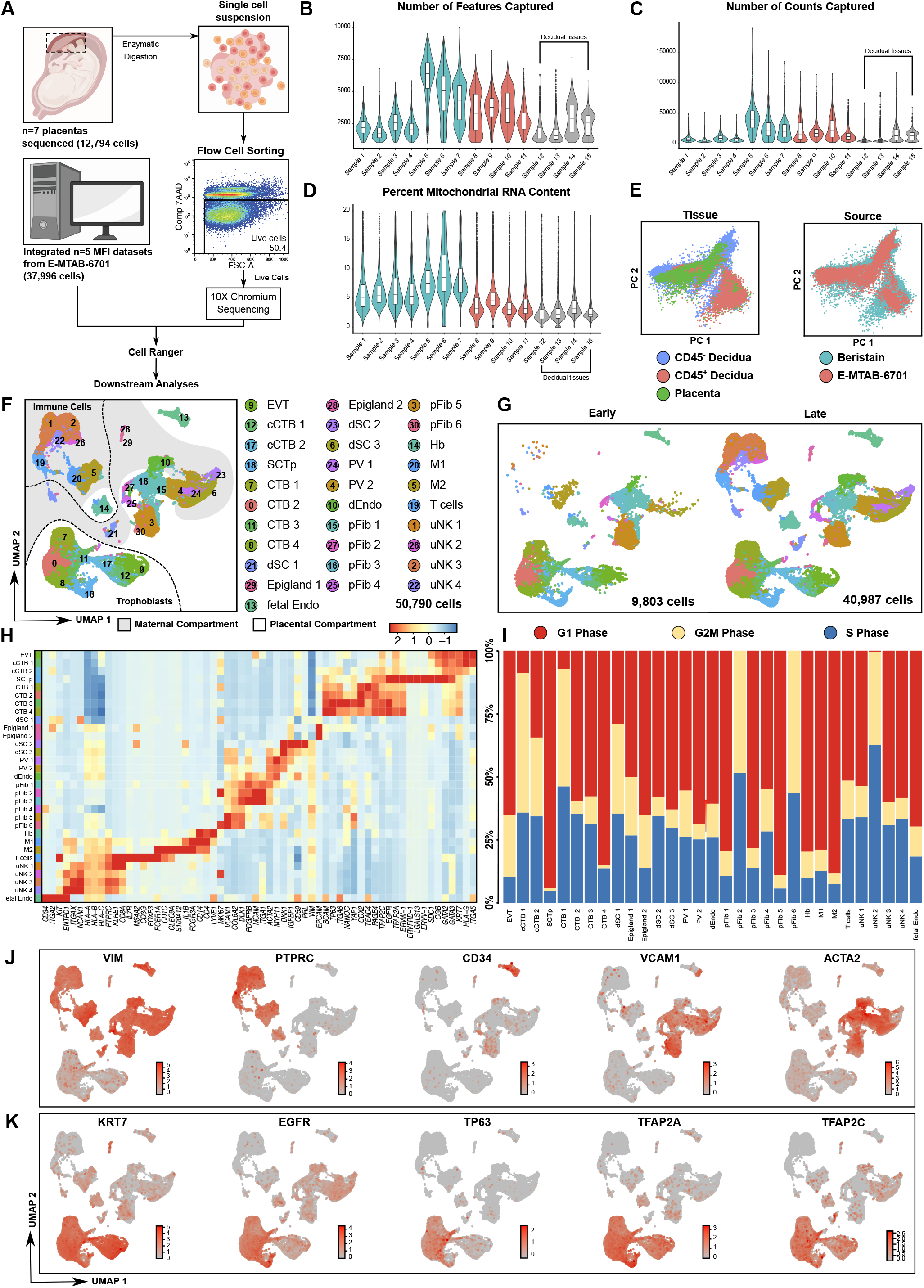
Sample quality control and single cell characterization of the first trimester human maternal-fetal interface. **(A)** Summary of workflow used for collection, processing, and single cell analysis of placental villous cells. MFI, Maternal-Fetal Interface, 7AAD: 7-Aminoactinomycin D, FSC-A: Forward Scatter Area. Violin plots showing the distribution of the raw number of **(B)** features, **(C)** the raw number of counts, and **(D)** the percent mitochondrial DNA content remaining post quality control and data pre-processing captured per cell during sequencing for each sample. Samples 1-6 (teal) represent the placental tissue samples sequenced by the Beristain lab, samples 7-11 (red) represent the placental tissue samples acquired from ArrayExpress, and samples 12-15 (grey) represent the decidual tissue samples acquired from ArrayExpress. **(E)** Scatter plot showing the distribution of captured cells across principal components 1 and 2 following dimensional reduction in principal component space. Cell positions are calculated based on cell embeddings calculated during dimensional reduction by principal component analysis. Cells are colour coordinated by source with teal representing those cells originating from placental tissue samples sequenced in this study and red representing those cells originating from placental and decidual tissue samples acquired from Vento-Tormo et al (ArrayExpress: E-MTAB-6701). **(F)** UMAP plot of 50,790 captured cells, demonstrating 31 distinct clusters identified within the first trimester human maternal fetal interface. Dashed lines separate maternal decidual immune cells from non-immune cells and placental trophoblasts. Cells are colour coordinated by identity and abbreviations are identical to those in Figure 1A. Epigland: epiglandular cell, fetal Endo: fetal endothelial cell, dEndo: decidual endothelial cell, dSC: decidual stromal cell PV: perivascular cell, pFib: placental fibroblast, Hb: fetal Hofbauer cell, M1: macrophage cell 1, M2: macrophage cell 2, uNK: uterine natural killer cell. **(G)** UMAP projection of the captured maternal fetal interface cells, stratified by gestational age. Early (left panel): 9803 early first trimester cells from each identified cluster captured between 6-7 weeks gestational age, before the onset of maternal placental vascularization. Late (right panel): 40,987 late first trimester cells from each identified cluster captured between 9-12 weeks gestational age, after the onset of maternal placental vascularization. Cell cluster names and colours correspond to those in Supplemental Figure 1E. **(H)** Heatmap showing the average expression of known immune, non-immune, and trophoblast gene markers, as well as proliferation gene markers in each identified cluster. Cell cluster names and colours correspond to those in Supplemental Figure 1E. **(I)** Stacked bar plot showing the percent distribution of cells in each cluster with gene signatures representing a cell in the gap 1 phase of proliferation, the gap 2 or mitotic phases of proliferation, or the synthesis phase of proliferation. Cell cluster names and colours correspond to those in Supplemental Figure 1E and abbreviations are identical to those in Figure 1F. **(J)** UMAP plots showing gene expression in red of a mesenchymal cell gene marker (*VIM*), an immune cell gene marker (*PTPRC*, encoding the CD45 protein), endothelial cell gene markers (*CD34* and *VCAM1*), and a fibroblast gene marker (*ACTA2*). **(K)** UMAP plots showing gene expression in red of known trophoblast gene markers used to subset and purify first trimester human trophoblasts from other cells within the maternal fetal interface.

**Supplemental Figure 2: 3D-rendering of UMAP-informed trophoblast clusters**

Chorionic villi scRNA-seq clusters showing state-specific colors corresponding to CTB1-4, SCTp, cCTB1, cCTB2, and EVT.

**Supplemental Figure 3:**
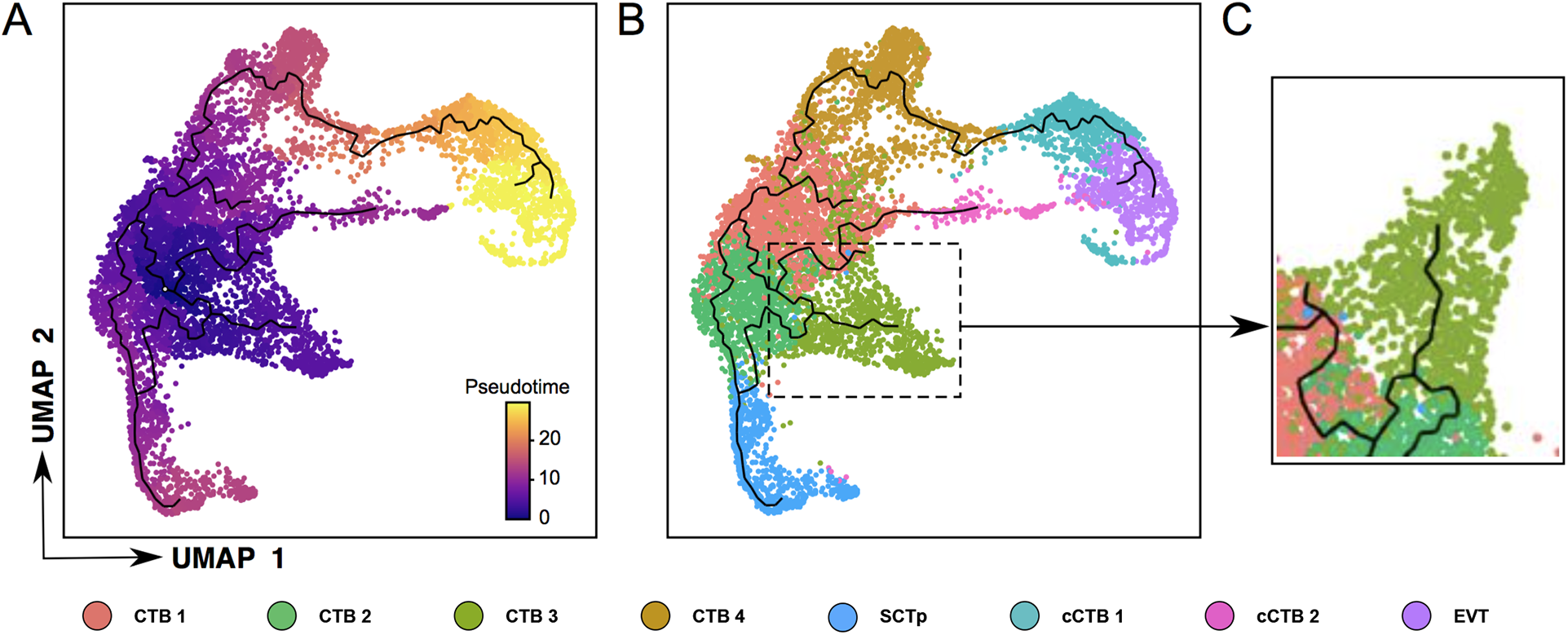
Monocle3 pseudotime quantification supports identification of CTB2 as the progenitor origin. **(A)** UMAP plot with constructed trajectory graph overlain. Cells are colour coordinated by relative pseudotime value. **(B)** UMAP plot with constructed trajectory graph overlain. Cells are colour coordinated by cluster identity. Cell cluster names and colours correspond to those in Figure 1C. **(C)** UMAP plot highlighting the constructed trajectory graph which describes the path taken as CTB state 2 progresses into CTB state 3. Cells are colour coordinated by cluster. Cluster names and colours correspond to those in Figure 1C.

**Supplemental Figure 4: 3D-rendering of UMAP-informed organoid trophoblast clusters** Trophoblast organoid scRNA-seq clusters showing state-specific colors corresponding to CTB1-3, SCTp, cCTB, EVT1, and EVT2.

**Supplemental Figure 5:**
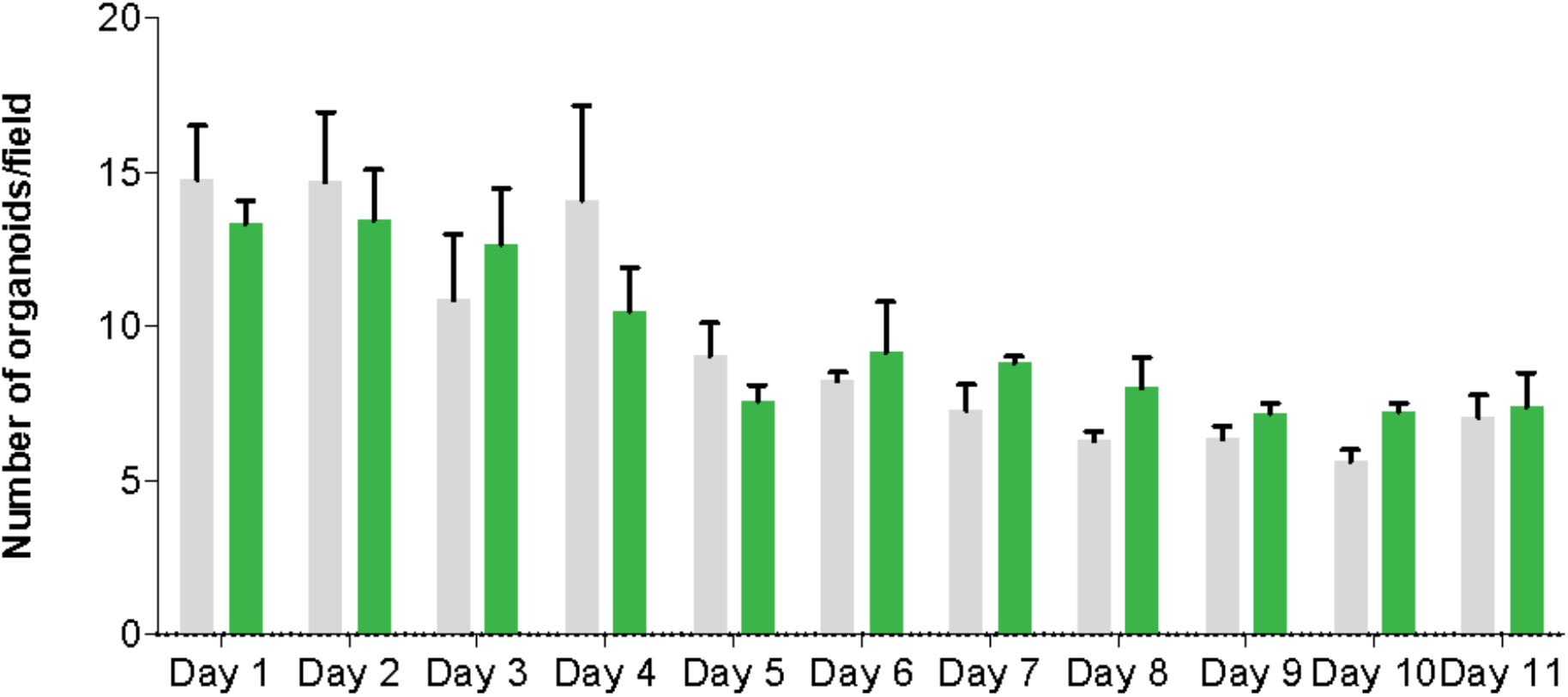
Organoid colonies in BCAM^hi^ and BCAM^lo^ sorted organoids. Bar graph shows the average of numbers of organoid colonies from BCAM^hi^ and BCAM^lo^ –sorted CT29 organoids cultured over 11 days. Grey bars = BCAM^lo^; Green bars = BCAM^hi^.

